# Spontaneous activity in peripheral sensory nerves – a systematic review

**DOI:** 10.1101/2023.06.13.544736

**Authors:** Dongchan Choi, George Goodwin, Edward B. Stevens, Nadia Soliman, Barbara Namer, Franziska Denk

**Affiliations:** Wolfson Centre for Age-Related Diseases, Guy’s Campus, King’s College London; Metrion Biosciences Ltd, Building 2 Granta Centre, Granta Park, Cambridge CB21 6AL; Imperial College London, Pain Research Group, Chelsea and Westminster Hospital, 369 Fulham Road, London, SW10 9NH; Research group Neuroscience of the Interdisziplinary Center for Clinical research, University Hospital of the RWTH Aachen, Aachen, Germany; Institute for Neurophysiology, University Hospital of the RWTH Aachen, Aachen, Germany

## Abstract

In the peripheral nervous system, spontaneous activity in sensory neurons is considered to be one of the two main drivers of chronic pain states, alongside neuronal sensitization. Despite this, the precise nature and timing of this spontaneous activity in neuropathic pain is not well-established.

Here, we have carried out a systematic search and data extraction of existing electrophysiological literature to shed light on which fibre types have been shown to maintain spontaneous activity and over what time frame. We examined both *in vivo* recordings of pre-clinical models of neuropathic pain, as well as microneurography recordings in humans.

Our analyses reveal that there is broad agreement on the presence of spontaneous activity in neuropathic pain conditions, even months after injury or years after onset of neuropathic symptoms in humans. However, due to the highly specialised nature of the electrophysiological methods used to measure spontaneous activity, there is also a high degree of variability and uncertainty around these results. Specifically, there are very few directly controlled experiments, with little directly comparable data between human and animals.

Given that spontaneous peripheral neuron activity is considered to be a key mechanistic feature of chronic pain conditions, it may be beneficial to conduct further experiments in this space.

## Introduction

Unlike cortical neurons, somatosensory afferents do not exhibit spontaneous activity under normal physiological conditions but only during pathological states. For example, one of the primary features of neuropathic pain in animal models is spontaneous activity in all major fibre types, including large myelinated Aβ, thinly myelinated Aδ and unmyelinated C fibres [14]. Similarly, in humans who live with neuropathy, spontaneous peripheral neuron activity can be measured using microneurography [13; 20; 26]. Based on these observations, numerous analgesic drug development strategies are aimed at reducing this abnormal pathological activity [31].

To inform these strategies, it is important to establish a consensus view of existing literature that goes beyond usual narrative summaries. We have conducted a review with systematic search and data extraction, a less biased method that can also provide insight into research quality [24].

We focused on experimental designs that can directly interrogate spontaneous neuron firing in whole organisms, i.e. *in vivo* electrophysiology and microneurography. Our first aim was to identify and collate all the relevant studies in this area into one, easily accessible spreadsheet. Our second aim was to use the data we extracted to answer a few simple, but crucial questions about spontaneous activity in neuropathy:

1. Which fibre types are mainly reported to be spontaneously active? Microneurography studies frequently report spontaneous activity in C fibres in neuropathy patients, e.g. [13; 19; 27]. Conversely, *in vivo* electrophysiology in animal models has traditionally pointed towards the importance of spontaneous activity in Aβ fibres, with Marshall Devor advocating that they are the main drivers of subsequent central sensitisation and pain [8].
2. How long does spontaneous activity persist? The tacit assumption of many in the field is that spontaneous activity is long-lasting and can drive ongoing pain. Indeed, in chronic neuropathy patients, spontaneous C-fibre firing can be observed which correlates with spontaneous pain [13]. The picture emerging from pre-clinical *in vivo* electrophysiology is much more mixed. Many report the bulk of activity within the first two weeks of nerve injury e.g. [22; 28], though others have argued that long-lasting activity emerges in unmyelinated fibres 3-4 weeks after an insult [16].
3. Are muscle afferents especially prone to sensitisation as opposed to skin afferents, as has been claimed [17]? If true, this could provide one potential mechanistic explanation for why deep muscle pain is a common complaint in chronic pain conditions.

The data we extracted supported the presence of ongoing spontaneous activity in both A and C fibres. However, there was a high level of heterogeneity and only very few, directly controlled studies. Microneurography studies were rare and spread across a divergent range of pain conditions. We conclude that the strength of available evidence in this area does not quite match the levels of confidence and ubiquity with which we all like to assert that spontaneous activity is one of the origins and main drivers of chronic pain.

## Methods

### Protocol Registration

We registered a systematic review protocol for our study on the Open Science Framework on 8 June 2020. However, as abstract screening and data extraction proceeded, it became clear that key papers had been missed using the search string that was originally registered, and that the literature was even more heterogeneous than initially anticipated. Significant deviations from the protocol therefore had to be undertaken to ensure inclusion of all the relevant literature. Moreover, data extraction was simplified to enable processing of the information within the time-frame of the project. Below contains a description of the methods that were eventually adopted.

### Systematic search

We conducted a systematic search of the existing literature using PubMed, to identify and collate evidence of spontaneous activity in sensory nerve fibres after peripheral injury. Abstracts were imported into EndNote for subsequent screening steps.

The search terms we initially used on the 8^th^ of June 2020 and logged as part of our systematic review protocol are presented as a string in Box 1. Medical Subject Headings (MeSH), i.e. specialised search strings designed for indexing, were used to exclude certain fields of research and limit the number of hits obtained. Once it became apparent that key papers were missing, another round of abstract screening and data extraction was undertaken. We added to already included papers with two additional search strings: that provided in Box 2 and the very broad string of “microneurography AND (spontaneous OR ongoing)”, which was designed to ensure that no relevant human microneurography papers would be missed. The second search was performed on 5^th^ of May 2021.

The main differences between the search strings in Box 1 and Box 2 were 1) the inclusion of ‘neuroma’ and ‘microsympathectomy’ before the first ‘AND’, 2) the inclusion of additional terms after the first ‘AND’, specifically ‘nerve impulse’, ‘microneurography’, ‘neuropathy’, ‘sympathectomy’, ‘nerve afferent’, ‘sympathetic efferent’, ‘electrophysiology AND pain’; and 3) the use of different MESH terms for exclusion, e.g. we added ‘brain diseases’ and removed ‘gene expression regulation’.

Our search strings and eventual data extraction relied on the terms ‘spontaneous, ‘ectopic and ‘ongoing’. We did not distinguish between the root causes of the spontaneous, ectopic or ongoing activity. For example, is the activity ‘’spontaneous’ in the sense of being generated endogenously by the neuron, i.e. as a result of membrane oscillations or changes in sodium channel function, or is it actually triggered by external factors, like chemical mediators released in the environment of injured neurons, and therefore, arguably not truly ‘spontaneous’ [2]. We did not distinguish between these within this review.

### Title and Abstract screening

Title and abstract screening was performed using the Systematic Review Facility (SyRF; http://syrf.org.uk/), an open-source systematic review platform [1]. Prior to screening, duplicate search results were removed by EndNote. If EndNote did not discard duplicates during the import stage, reviewers removed any remaining duplicate articles. The remaining unique references were then screened for eligibility against the below inclusion/exclusion criteria using the title and abstract text. If a decision could not be reached on this information alone, the full text of an article was accessed in a second step. Each reference was evaluated by two independent reviewers, specifically, DC & FD. Disagreements were resolved through joint discussion, erring on the side of inclusion in the first instance.

#### Box 1.

##### Search string registered in the original protocol, used for the first round of data collection.

*Note that the PubMed database does not distinguish between American and English spelling, i*.*e. the term ‘fibre’ will find both ‘fibre’ and ‘fibre’*.

(Spontaneous discharge* OR spontaneous firing* OR Spontaneous activit* OR ongoing discharge* OR ongoing firing* OR ongoing activit* OR ectopic discharge* OR ectopic firing* OR ectopic activit* OR ectopi*) AND ((”Sensory neuron*”) OR (”A fibre*”) OR (”C fibre*”) OR (”neuropathic pain”) OR (”nerve injur*”) OR (”axotom*”) OR (”Phantom pain”) OR (”neuroma*”) OR (”severed nerve*”)) NOT REVIEW NOT (”Cardiovascular Diseases” [mesh]) NOT (”Electroencephalography” [mesh]) NOT (”Memory Disorders”[MESH]) NOT (”intellectual disability”[mesh]) NOT (”consciousness disorders”[mesh]) NOT (”cochlear nerve” [MESH]) NOT (”Vestibulocochlear Nerve” [mesh]) NOT (”Optic Nerve”[mesh]) NOT (”oculomotor muscles” [mesh]) NOT (”tomography”[mesh]) NOT (”Electroencephalography” [mesh]) NOT (”computational biology”[mesh]) NOT (”Receptors, Odorant”[Mesh]) NOT (”Gene Expression Regulation”[Mesh]) NOT (”Congenital, Hereditary, and Neonatal Diseases and Abnormalities” [mesh]) NOT (”Respiratory Tract Diseases” [mesh]) NOT (”olfactory bulb” [mesh]) NOT (”intestine, small”[MESH]) NOT (”invertebrates”[MeSH]) NOT (”fishes” [MESH]) NOT (”Herpesviridae”[mesh]) NOT (”Ear Diseases” [MESH]) NOT (“Magnetoencephalography”[Mesh]) NOT (”Joint Diseases”[Mesh]) NOT (”Information Science”[Mesh]) NOT (”Microglia”[Mesh])

#### Box 2.

##### Second search string used to for the second round of data collection.

(Spontaneous discharge* OR spontaneous firing* OR Spontaneous activit* OR ongoing discharge* OR ongoing firing* OR ongoing activit* OR ectopic discharge* OR ectopic firing* OR ectopic activit* OR ectopi* OR neuroma* OR microsympathectomy) AND ((”Sensory neuron*”) OR (”A fibre*”) OR (”C fibre*”) OR (”neuropathic pain”) OR (”nerve impulse*”) OR (”nerve injur*”) OR (”axotom*”) OR (”electrophysiology” AND “pain”) OR (”microneurography”) OR (”neuropathy”) OR (”Phantom pain”) OR (”Sympathectomy”) OR (”nerve afferent*”) OR (”sympathetic efferent*”)) NOT REVIEW NOT (”BRAIN”[mesh]) NOT (”intestine, small”[MESH]) NOT (”invertebrates”[MeSH]) NOT (”fishes” [MESH]) NOT (”Herpesviridae”[mesh]) NOT (”Ear Diseases” [MESH]) NOT (”brain diseases” [MESH]) NOT (”Ocular Motility Disorders”[mesh]) NOT (”Vestibulocochlear Nerve Diseases”[mesh]) NOT (”eye diseases”[mesh]) NOT (”Cardiovascular Diseases”[mesh]) NOT (”cardiovascular physiological phenomena”[MESH]) NOT (”Memory Disorders”[MESH]) NOT (”intellectual disability”[mesh]) NOT (”consciousness disorders”[mesh]) NOT (”mental processes”[mesh]) NOT (”Stomatognathic Diseases” [mesh]) NOT (”cochlear nerve” [MESH]) NOT (”Vestibulocochlear Nerve” [mesh]) NOT (”Optic Nerve”[mesh]) NOT (”oculomotor muscles” [mesh]) NOT (”tomography” [mesh]) NOT (”Electroencephalography” [mesh]) NOT (”computational biology”[mesh]) NOT (”psychophysiology”[mesh]) NOT (”neoplasms”[mesh]) NOT (”Congenital, Hereditary, and Neonatal Diseases and Abnormalities” [mesh]) NOT (”Respiratory Tract Diseases” [mesh])

### Inclusion and Exclusion Criteria

During abstract screening of articles concerning non-human data, we included all primary articles that appeared to present any electrophysiology data of spontaneous peripheral neuron activity resulting from nerve injury. We excluded review articles or articles that only showed schematic drawings of electrophysiology recordings rather than actual data.References to textbooks were excluded. During subsequent data extraction, we included only full-text articles, written in English, that performed *in vivo* electrophysiological recordings of peripheral sensory nerves in models of neuropathic pain. We did not include articles examining spontaneous activity after inflammation or in diabetic models. Arguably, diabetes could cause neuropathy, but there were only four papers which attempted to study spontaneous activity in animal models of diabetes, three of which used the streptozotocin model which is not considered to have particularly good face validity [11].

During abstract screening of articles concerning human data, we included only original articles, which appeared to contain microneurography recordings. Here, data of patients with any sort of chronic pain were included. Reviews were excluded. For data extraction, the full text of the article had to be available in English.

Qualitative data collection (e.g. species, sex, type of neuropathic pain model, human disease/pain type) was performed on all the articles included for full data extraction. However, quantitative data, e.g. on n-numbers and spontaneous activity, was only obtained if the papers also provided raw data in a format that permitted us to calculate the percentage of spontaneously active fibres recorded in a particular condition.

### Data extraction and management

Once screened, all references and their corresponding abstracts returned from the searches were downloaded from SyRF and compiled into an Excel Database (**Supplementary Table**, [available on OSF](https://osf.io/6wpnq/?view_only=12dad701949b4ea88c2a8c3b0b50fa7c)). This database contains the list of included articles, and specifically information on title, authors, journal, and year of publication. Two reviewers (DC & FD) independently went through each item, decided on full text inclusion, and if suitable, extracted data. Given how different *in vivo* electrophysiology and microneurography are as techniques, different parameters were extracted for non-human and human data, as described in the following:

#### For non-human data

reasons for exclusion (if applicable), study purpose, animal species, animal sex (where “unclear” was used to denote papers in which no information was provided), injury type (e.g. sciatic nerve ligation) and model type. Options for the latter included “non-regenerating” (e.g. nerve ligation), “regenerating” (e.g. nerve crush), “non-traumatic” (e.g. chemotherapy-induced neuropathy), “other” (e.g. rhizotomy). We also collected information on the type of nerve that was being injured (e.g. “sciatic”), as well as whether the nerve in question contained cutaneous afferents, muscle afferents or a mixture of both. Data were extracted on fibre type (e.g. “A fibre”), how fibre type was determined (e.g. using conduction velocity) and how much time after injury the recordings were conducted: acute (= less than 24h), less than 8 days, weeks (i.e. 8-21 days), chronic (i.e. more than 21 days). We noted where the activity was recorded from, e.g. in teased L5 dorsal root, and how many individual animals and/or filaments were used. The number and/or percentage of spontaneously active fibres recorded per group was extracted. Finally, we also scored the ‘level of bias’ of each article, where bias refers to how likely the recorded incidence of spontaneous activity would have been impacted by how the data were acquired. Specifically, C fibres are known to sensitise and start firing spontaneously in response to sustained activation, e.g. as a result of repetitive receptive field testing [5]. We ranked articles based on our interpretation of how likely recording bias would have resulted, given the authors’ description of their experiments. Articles were scored ‘low bias’ when authors specifically mentioned taking care not to sensitise fibres or clearly avoided repetitive stimulation of fibres prior to recording of spontaneous activity; articles were scored ‘medium bias’ when some initial receptive field testing or electrical stimulation had taken place; and finally, articles were scored ‘high bias’, if the authors specifically focused their analyses on those fibres which were spontaneously active or conducted extensive receptive field stimulation.

*For human data*, we recorded the overall purpose, the sex of the participants, the injury or disease, the number of patients from whom spontaneous activity data was recorded, and whether they experienced painful or non-painful neuropathy or whether they were healthy controls. Where available, we also recorded how many individuals were reported to experience spontaneous activity, whether this was associated with spontaneous pain, and whether any quantitative sensory testing was performed. Finally, we extracted the number of filaments that were being recorded from and how many of them were spontaneously active, again grouped into painful and non-painful neuropathy groups vs. healthy controls. Acquisition bias was not considered for microneurography recordings.

In all instances, data extraction proceeded by experiment, with some articles containing several experiments, and thus several pieces of information, each provided with their own unique experiment number. An experiment was defined based on whether it generated its own instance of spontaneous activity data, e.g. spontaneous activity measured at a different time point, at a different site, or under a different condition was logged separately.

Since control data were lacking in most experiments, we chose to plot the percentage of spontaneously active fibres that were being reported. For statistics, we performed non-parametric tests, but given that this was not a hypothesis-testing endeavour and given the highly heterogeneous nature of the underlying data, we invite the reader to focus on the median and quartile ranges provided, rather than on probability statistics [29].

## Results

A study flow diagram starting from database search to extraction is illustrated in **Figure 1**. We obtained 1170, 1218 and 110 results on Pubmed using the initial, second and third search strings, respectively. After removal of 565 duplicates, primary screens based on title and abstract reduced the number of articles to 336 unique articles on non-human data and 57 articles on human microneurography data.

**Figure 1.**
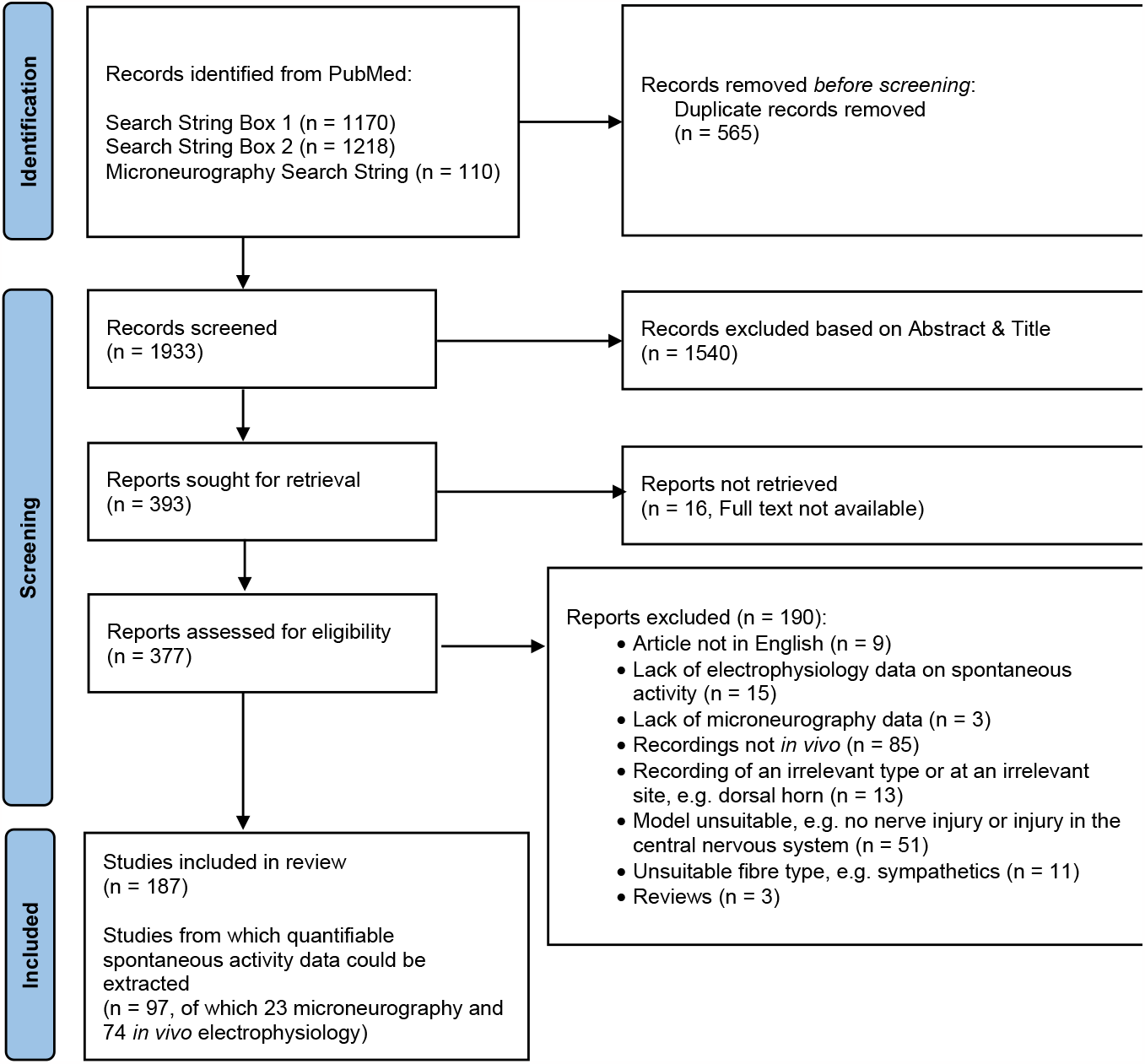
Systematic search flow chart.

During full text screening, we excluded 190 articles for a variety of reasons specified in **Figure 1** and the ‘reason_meta’ column in **Supplementary Table 1**. Of the 187 articles that were subsequently included, 74 *in vivo* electrophysiology and 23 microneurography articles were taken for full quantitative data extraction, while the remaining 90 were processed qualitatively. For the former, we extracted spontaneous activity data from individual experiments. For the latter, we extracted other information, e.g. on species, gender, condition and/or nerve injury models.

**Table 1.**
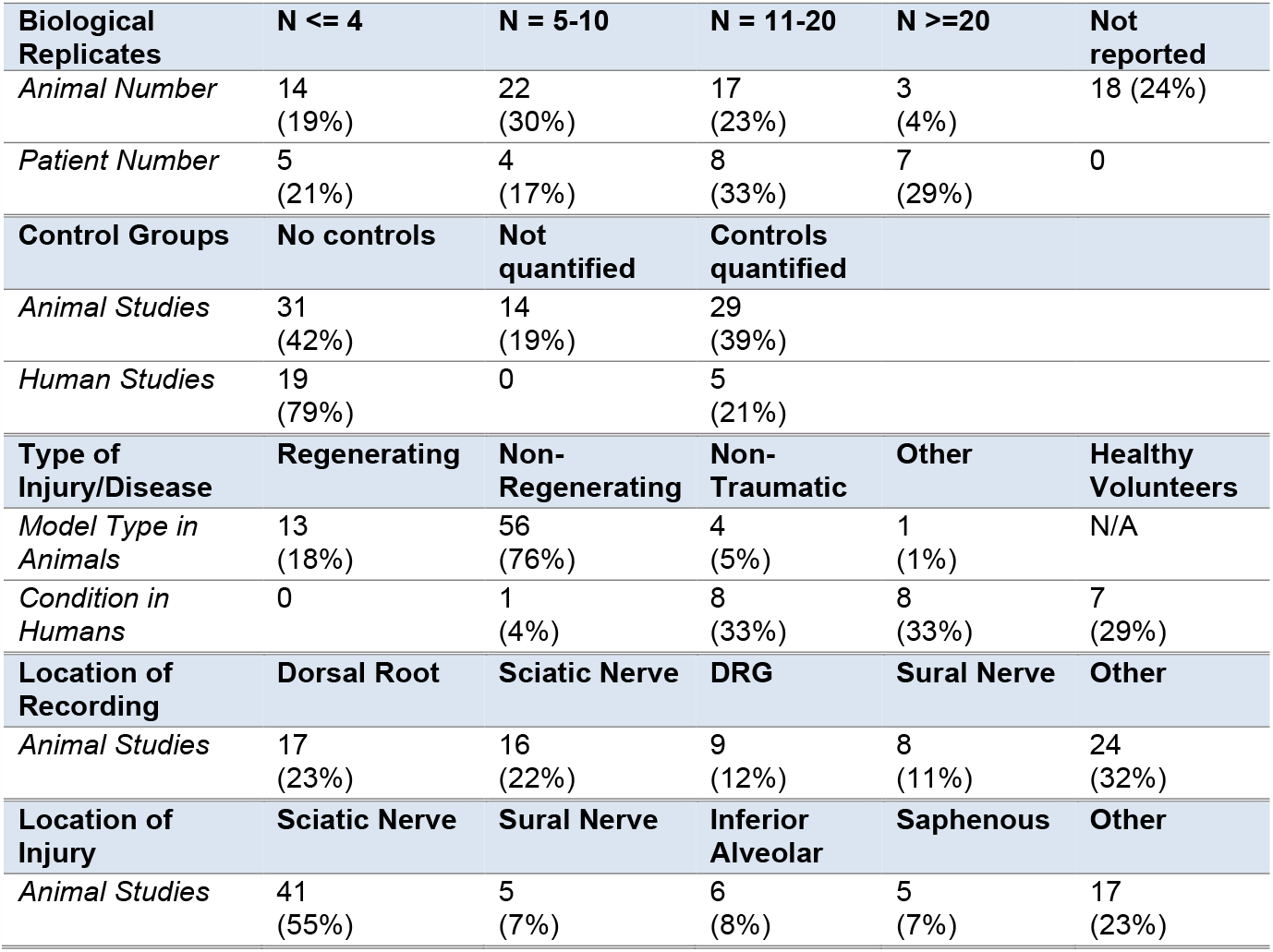
High heterogeneity between the studies with quantifiable data on spontaneous activity. The 97 studies we included for full quantification varied greatly in terms of their experimental design, with varying n numbers, injury/disease type and location. Many of them also failed to provide recordings from control groups. NB: The table includes information on two separate experiments from one of the n = 23 human studies, as they were conducted on two patients with divergent syndromes (study 250 in Supplementary Table, on one individual with fibromyalgia and one with small fibre neuropathy).

### Characterisation of included studies

Of the non-human experiments, we unsurprisingly found that the majority of recordings were performed in rats (65%, n = 122), while the remainder were performed in ferret (5%, n = 10), cat (4%, n = 8), mouse (3%, n = 6) and monkey (1%, n =1). 21% of the studies we included (n = 40) were human microneurography studies (**Figure 2**).

**Figure 2.**
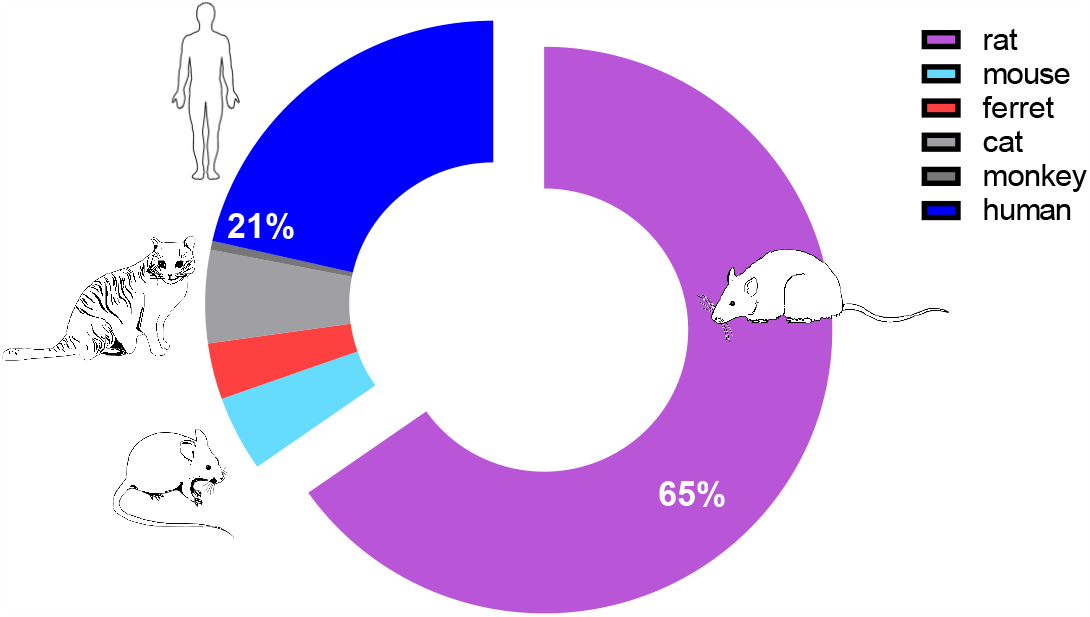
Rat and human data make up the majority of the dataset.

Among the non-human experiments, most studies were performed in male animals (62.6%, n = 92), while 12.2% used either only female or both sexes. The remaining 12.9% did not provide any information on sex. In human microneurography studies, the picture was reversed, with 67.5% of the work (n = 27) using both genders, 22.5% only females (n = 9), 7.5% only males (n = 3) and only 1 study not specifying (**Figure 3**).

**Figure 3.**
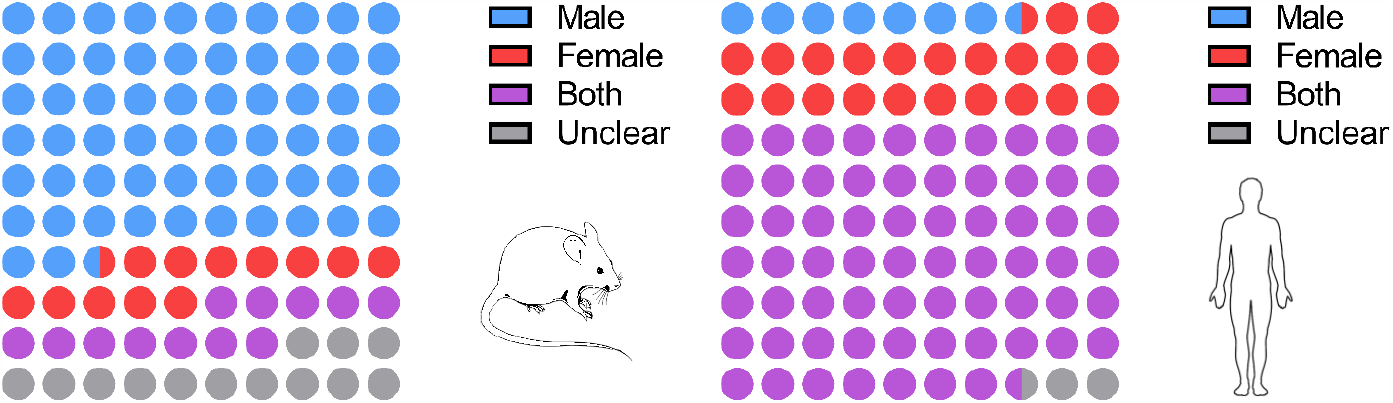
The majority of data collected from non-human studies (n = 147) were male-only, while human studies (n = 40) included both sexes. ‘Unclear’ was used as a category in cases where no information on sex or gender was provided.

### High variability in experimental design and reporting practises between studies

Among the articles we studied there was significant heterogeneity in what was being reported. For example, nearly half of all articles we included (73 animals and 17 human studies, 48% of the total 187) could not be used for quantitative analysis. Most commonly, they failed to report the total number of fibres measured or the raw number of spontaneously active fibres, making it impossible to calculate the proportion of neurons with spontaneous firing. Also, many studies were designed to only record from spontaneously active units.

Of the 97 articles that could be used for full quantification, there was high heterogeneity among animal studies in terms of recording location and the site of injury (**Table 1**). Moreover, most studies in animals were on non-regenerating traumatic nerve injury models (76%), while human microneurography studies were relatively evenly split between recordings from healthy volunteers, those living with chronic primary pain conditions like fibromyalgia and those living with non-traumatic neuropathy e.g. as a result of diabetes or chemotherapy.

Another major source of heterogeneity was that many articles did not include a control group, but simply measured the percentage of spontaneously active neurons in a pain condition. This was true for both *in vivo* electrophysiology on animal models, and even more so for microneurography data. Thus, 79% of microneurography studies did not include concurrent recordings from more than one condition, e.g. a given article might only include data from neuropathy patients or only use historically generated control data. Similarly, 42% of animal studies did not include any control condition, whether it be sham, naïve or vehicle-treated animals. There were also many studies that used the same control group for a series of experimental comparisons or did not provide quantifiable data on the level of spontaneous activity in their controls. In total, there were therefore only 29 articles that allowed for direct comparison between injury and control groups (39%).

Finally, in terms of reporting practises, *in vivo* electrophysiology studies, as a rule, considered individual fibres or filaments as the unit of measurement used in discussions and subsequent statistical analyses. The total animals that were used to generate the filament data would either not be reported at all (24% of cases) or reported, but on a summary level. For example, an article might cite that they used n = 20 rats in total, but it would not be made clear which filaments were recorded in which animal. Much information on inter-animal variability has therefore been lost from the literature examining spontaneous activity.

### Quantitative spontaneous activity data

Within the 74 articles containing quantitative data from non-human models, we extracted 126 experiments conducted on A fibres and 88 experiments on nociceptors, including C and Aδ fibres. For 21 experiments, the fibre type was not reported. The included studies used a bewildering number of different neuropathic pain models, requiring 20 different terms to allow full categorisation, including spinal nerve ligation, chronic constriction injury, nerve crush, transection, nerve ligation (of the peripheral branch), neuroma and chemotherapy-induced neuropathy (see column ‘injury_type’ in **Supplementary Table**).

Classifying these various neuropathic pain models into four broader categories showed that 76% of experiments were conducted on non-regenerative models (e.g. nerve ligation), 18% on regenerative models (e.g. nerve crush), 4% on non-traumatic neuropathic pain models (induced by chemotherapy, neuritis or ischemia), and 1% on other injury models including rhizotomy, dorsal column transection and disc rupture (**Table 1 & Supplementary Table**, column ‘Model Type’). All recorded model categories showed an increase in the median percentage of spontaneously active fibres compared to controls (**Figure 4 & Table 2**), with the difference reaching statistical significance for non-regenerating and non-traumatic categories (Mann-Whitney tests, q = 0.00001 and q = 0.0002, respectively). In the regenerating group, the mean rank of the injury group was higher (18.34) compared to the control group (12.81), but this difference was not statistically significant when examined with a Mann-Whitney t-test.

**Table 2.** Medians with lower & upper quartiles of the values plotted in Figures 4 & 5. Numbers represent the percentage of spontaneous activity.

| **All experimental models** | **Control Median** | **Control Quartiles** | **Injury Median** | **Injury Quartiles** |
| --- | --- | --- | --- | --- |
| Non-Regenerating | 3.0 | 0 - 5 | 8.0 | 4 - 13.8 |
| Regenerating | 0.5 | 0 - 9.8 | 7.0 | 0 - 19.5 |
| Non-Traumatic | 1.5 | 0 - 4.8 | 10.5 | 4.5 - 21 |
| Other | 11.0 |  | 10.0 | 1 - 25 |

| **Experiments with internal controls only** | **Control Median** | **Control Quartiles** | **Injury Median** | **Injury Quartiles** |
| --- | --- | --- | --- | --- |
| *Non-Regenerating* | 3.0 | 0 - 5 | 13.0 | 8 - 22.8 |
| *Regenerating* | 0.0 | 0 - 3 | 7.0 | 0 - 10 |
| *Non-Traumatic* | 1.5 | 0 - 4.8 | 13.5 | 7 - 23 |
| *Other* | 11.0 |  | 25.0 |  |

**Figure 4.**
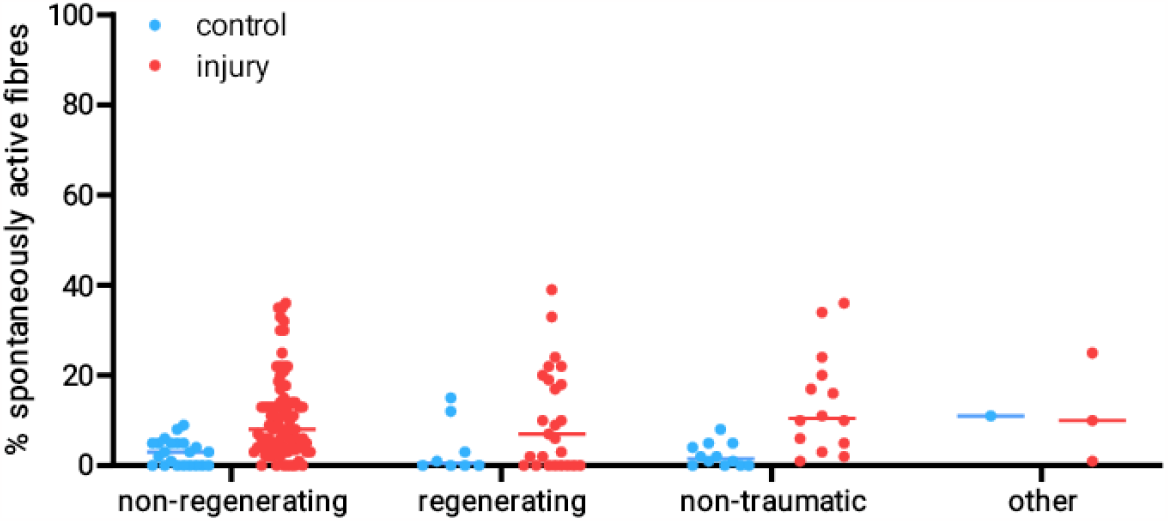
The percentage of spontaneously active fibres was increased across all experimental categories. Plotted here is the percentage of spontaneously active fibres recorded at any time post injury in experiments categorised as “low bias”. Experiments which injured pure muscle afferents were excluded. Each dot is an experiment, and the lines represent the medians for each group. non-regenerating: n = 22 for control, n = 88 for injury; regenerating: n = 8 for control, n = 25 for injury; non-traumatic (neuritis & chemotherapy-induced neuropathy due to oxaliplatin, paclitaxel & vincristine): n = 12 for control, n = 14 for injury; other (rhizotomy & dorsal column transection): n = 1 for control, n = 3 for injury.

It is important to bear in mind that the controls in **Figure 4** are not study-matched, so the results will be prone to batch effects, and only the largest effect sizes will be detectable. If only studies with their own control condition are considered, the number of experiments eligible for inclusion drop from 130 down to 42 (**Figure 5**). The overall conclusions remain unchanged though, with the difference between control and neuropathy reaching statistical significance for non-regenerating and non-traumatic categories (**Table 2**, Mann-Whitney tests, q = 0.00001 and q = 0.000044, respectively). In the regenerating group, the effect was once again less clear, though the mean rank of the injury group was still higher (8.43) compared to the control group (6.58).

**Figure 5.**
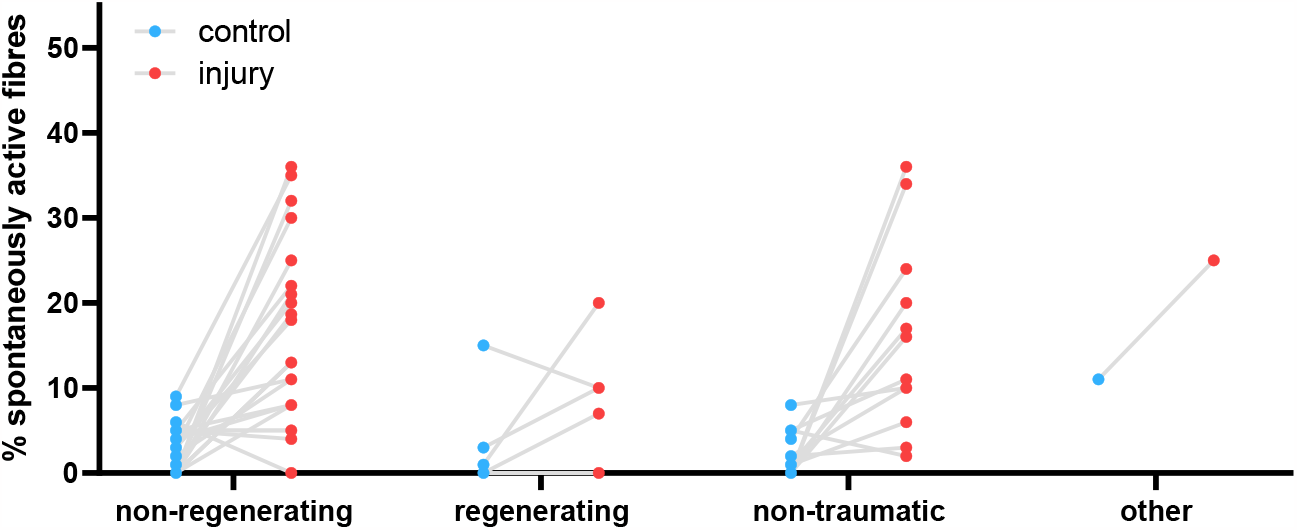
The percentage of spontaneously active fibres considering only studies with their own internal controls. Plotted here is the percentage of spontaneously active fibres recorded at any time post injury in experiments categorised as “low bias” which included their own controls. Experiments which injured pure muscle afferents were excluded. Each dot is an experiment, with the lines connecting control and neuropathy conditions. non-regenerating: n = 22; regenerating: n = 7; non-traumatic (neuritis & chemotherapy-induced neuropathy due to oxaliplatin, paclitaxel & vincristine): n = 12; other (L5 ventral rhizotomy): n = 1.

In humans, of those studies included for full quantitative data extraction, there were 5 experiments on A fibres, 22 on C nociceptors and 1 that was unclear. 7 articles included individuals with chronic primary pain (e.g. complex regional pain syndrome or fibromyalgia), 9 included those with neuropathies of some kind, e.g. as a result of diabetes, 7 included healthy volunteers, and one study was performed on patients with pain as a result of tooth decay. The proportion of spontaneously active C fibres was higher in those individuals living with painful neuropathy, compared to those living with non-painful neuropathies and healthy controls **(Figure 6)**.

**Figure 6.**
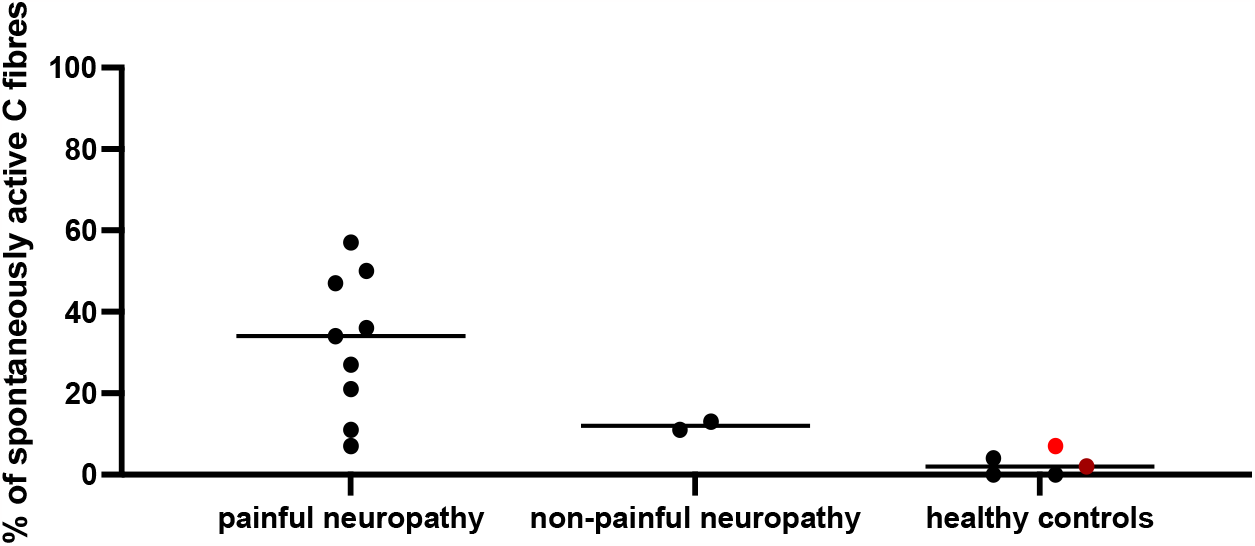
The percentage of spontaneously active C fibres was higher in those individuals living with painful neuropathy. Each dot is data derived from an individual experiment. Median values indicated by the line are listed in the following (confidence intervals and n numbers provided in brackets): painful neuropathy: 34 (11-50, n = 9); non-painful neuropathy: 12 (11-13, n = 2); healthy controls: 2 (0-7, n = 5). The dot in red is recorded from muscle C fibres, the one in dark red from aging healthy volunteers.

A Kruskal-Wallis test between the three groups revealed a statistically significant difference, specifically between painful neuropathy groups and healthy controls at adj. p = 0.0028. Only two of the studies plotted in **Figure 6** had their own internal controls [13; 21], with one of them reporting increased firing in painful neuropathy (27% of fibres vs. 13% of fibres [13]), while the other one only reported increased firing in neuropathy vs. control, regardless of pain status (4% of fibres in healthy controls vs. 7% and 11% in painful vs. non-painful neuropathy, respectively [21].

Including only studies with their own concurrent control groups in the same article, we can extract data from only 5 studies on painful peripheral neuropathy, fibromyalgia and pain due to tooth decay. Spontaneous activity was significantly increased in pain conditions compared to non-painful controls, as summarised in **Figure 7** (Mann-Whitney two-tailed t-tests, p = 0.0159).

**Figure 7.**
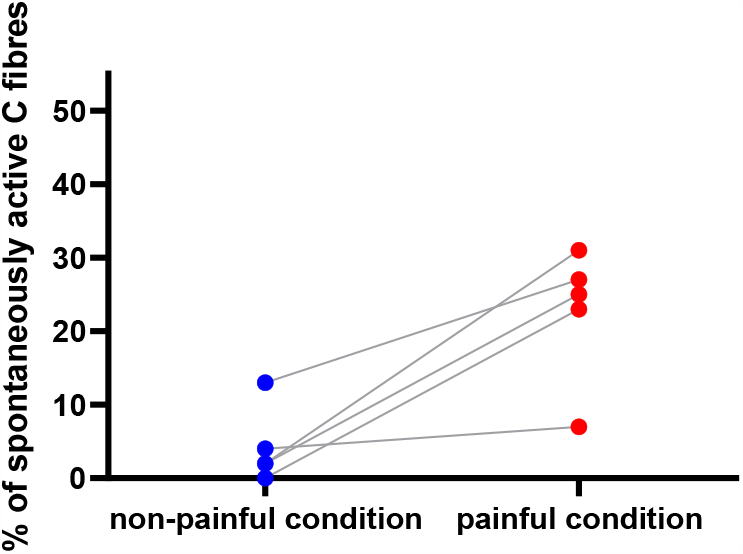
The percentage of spontaneously active C fibres from human microneurography studies with concurrent control recordings. Each dot is data derived from an individual experiment. The line connects recordings made in control vs. painful conditions. Data derive from five articles (2 × studying fibromyalgia patients, 2× peripheral neuropathy patients and 1× studying tooth decay): article numbers 32, 244, 246, 250 and 254 in **Supplementary Table**).

### Impact of bias, fibre type and time on spontaneous activity

To answer the question of how bias in the recording setup affected the percent of spontaneous activity recorded in the different fibre types, we split our non-human dataset according to our low, medium and high bias categories. We limited our analysis to non-regenerating traumatic neuropathic models and examined either A fibres (**Figure 8**) or Aδ and C fibres (**Figure 9**). The latter group are predominantly nociceptors, however, not all papers confirmed responses in the noxious range with receptive field stimulation. Moreover, much of the early literature did not further classify A fibres into Aβ and Aδ categories. Thus, the data in **Figure 8** may include activity from some nociceptive Aδ fibres.

**Figure 8.**
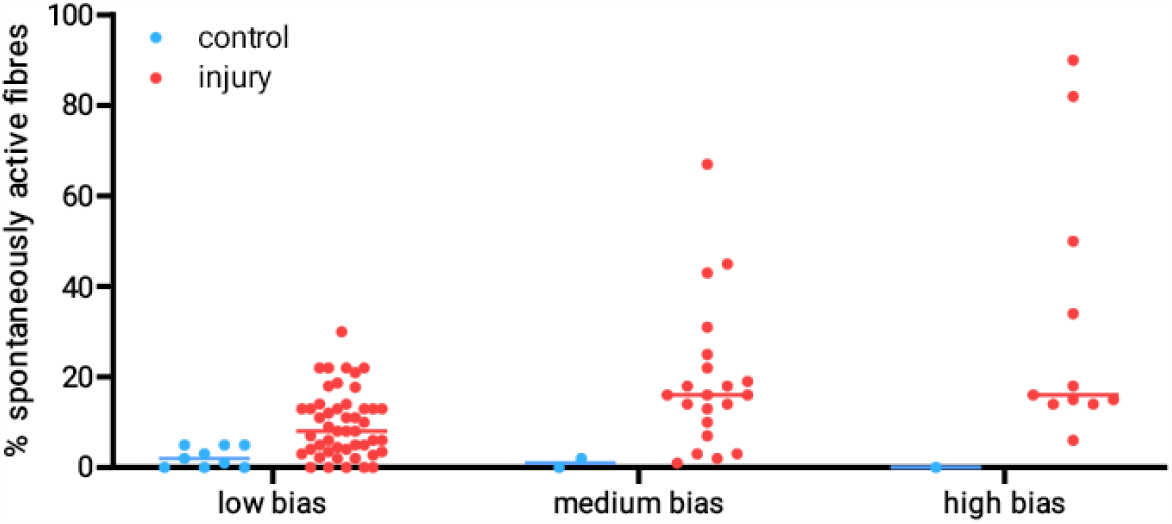
A fibres show increased spontaneous activity after injury when compared to control recordings. The data included in the plot are derived from experiments on A fibres recorded after injury of either mixed or cutaneous nerves in non-regenerative models. The line indicates the median values. The control group includes data from naïve, vehicle-treated and sham-injured animals. N numbers for control & injury groups: low bias n=9 & 48; medium bias n=2 & 21; high bias n= 1 & 11.

**Figure 9.**
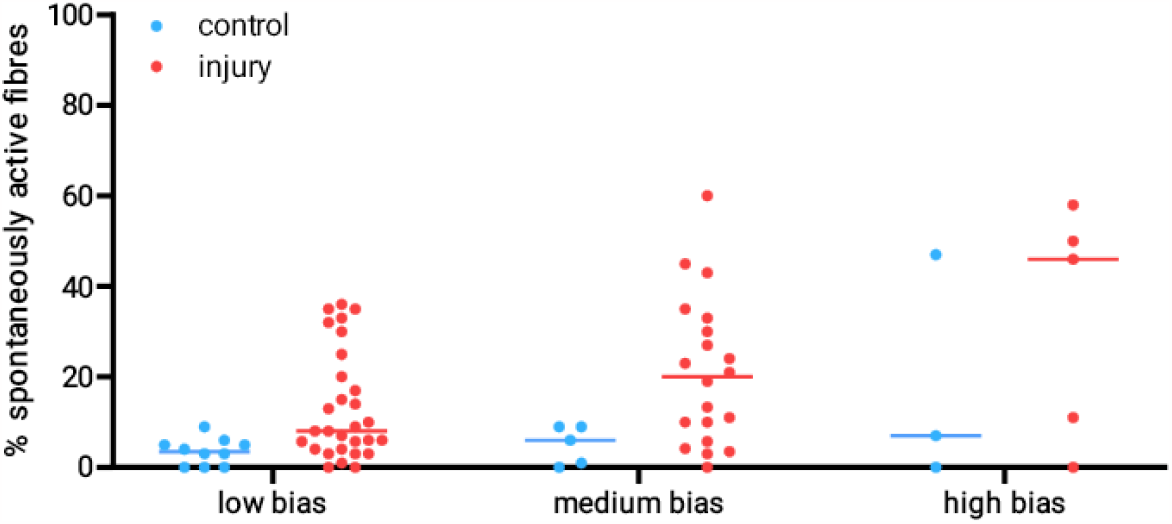
Putative nociceptors show spontaneous activity after injury when compared to control recordings. The data included in the plot are derived from experiments on C fibre and Aδ fibres recorded after injury of either mixed or cutaneous nerves in non-regenerative models. The line indicates the median values. The control group includes data from naïve, vehicle-treated and sham-injured animals. N numbers for control & injury groups: low bias n=10 & 29; medium bias n=5 & 20; high bias n= 3 & 5.

As expected, the percentage of spontaneously active fibres was higher in the medium and high bias groups, and this effect was particularly pronounced for Aδ and C fibres – putative nociceptors and therefore prone to sensitisation (**Table 3**). Accordingly, a mixed effects model revealed a main effect of injury for both A fibre and nociceptor experiments, but only a main effect of bias for nociceptor experiments: fixed effect of *injury* (A fibres): F(1, 86) = 9.885, p = 0.0023; fixed effect of *injury* (nociceptors): F(1,66) = 9.159, p = 0.0035; fixed effect of *bias* (nociceptors): F(1, 32) = 4.407, p = 0.049.

**Table 3.**
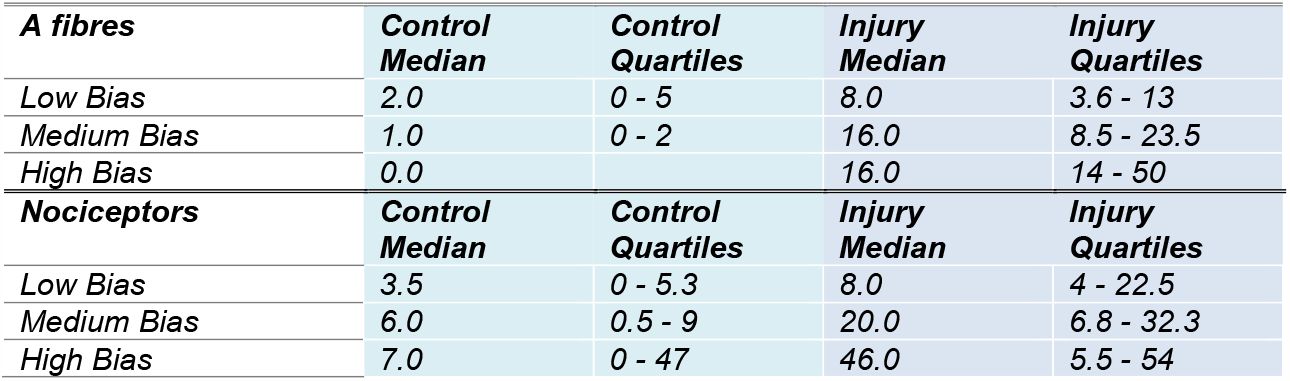
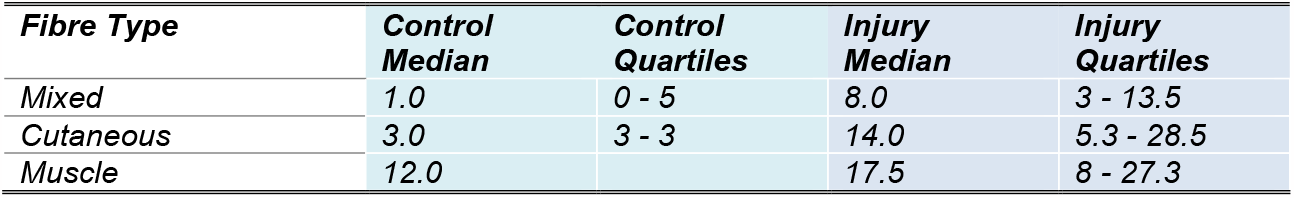
Medians with lower & upper quartiles of the values plotted in Figures 8 - 10. Numbers represent the percentage of spontaneous activity.

To investigate whether muscle fibres indeed display more spontaneous activity, as is usually claimed in the narrative literature, we split low bias experiments conducted on regenerative and non-regenerative models according to the type of nerve that had been injured: purely cutaneous (e.g. sural), purely muscle (e.g. gastrocnemius) or mixed (**Figure 10**). Numerically, the median of the percent spontaneous activity recorded after injury in individual experiments was indeed marginally larger in muscle nerves (median: 17.5, n = 12) compared to cutaneous (14.0, n = 12) and mixed nerves (8.0, n = 101), see also **Table 3**. Moreover, a Kruskal-Wallis test showed that this difference was significant comparing mixed vs. muscle nerves after injury (adj. p value = 0.029), while the difference between mixed vs. cutaneous nerves was not (adj. p value = 0.23).

**Figure 10.**
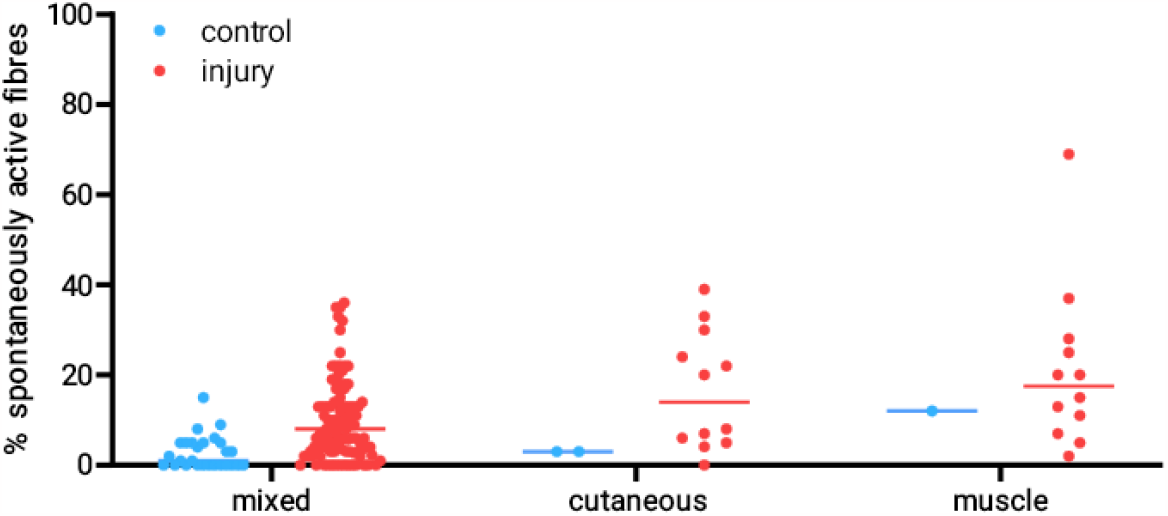
The percentage of spontaneously active fibres was higher after injury in muscle rather than cutaneous and mixed nerves. Each dot is data derived from an individual experiment. Only low bias experiments were included. The line indicates the median values. The control group includes data from naïve, vehicle-treated and sham-injured animals. N numbers for control & injury groups: mixed nerves n = 27 & 101; cutaneous nerves n=2 & 12; muscle nerves n= 1 & 12.

Finally, we split our data according to the amount of time that had elapsed after injury (**Figure 11A**). The vast majority of control experiments recorded less than 10% spontaneous activity. After nerve damage, spontaneous activity was observed in all fibre types, although for nociceptors, it was most prominently reported less than 8 days after injury. This may be partly due to experiments on C and Aδ nociceptors becoming less common at later time points. Pooling across fibre types, a Kruskal-Wallis test revealed a significant increase in spontaneous activity after injury for three of the groups compared to the control (**Figure 11B**): less than 8 days (adj. p < 0.0001), between 8 days and 3 weeks (adj. p = 0.013) and more than 3 weeks (adj. p = 0.031). See **Table 4** for medians and quartile ranges.

**Table 4.** Medians with lower & upper quartiles of the values plotted in Figures 11B & 12. Numbers represent the percentage of spontaneous activity.

| Condition | Median | Quartiles |  |  |
| --- | --- | --- | --- | --- |
| Control | 2.0 | 0 - 5 |  |  |
| Acute Injury | 5.0 | 0 - 13 |  |  |
| Injury <8 days | 13.0 | 6 - 22 |  |  |
| Injury after 1-3 weeks | 6.0 | 0 - 15.9 |  |  |
| Injury after 3 wks+ | 6.0 | 3 - 9.3 |  |  |
| Time Point | Control Median | Control Quartiles | Injury Median | Injury Quartiles |
| Acute | 5.0 |  | 5.0 | 4 - 22 |
| <8 days | 1.5 | 0 - 4.3 | 15.5 | 8 - 26.3 |
| Weeks | 0.0 | 0 - 0.8 | 0.0 | 0 - 15 |
| 3 wks+ | 8.0 | 1.5 - 13.5 | 10.0 | 9 - 13 |

**Figure 11.**
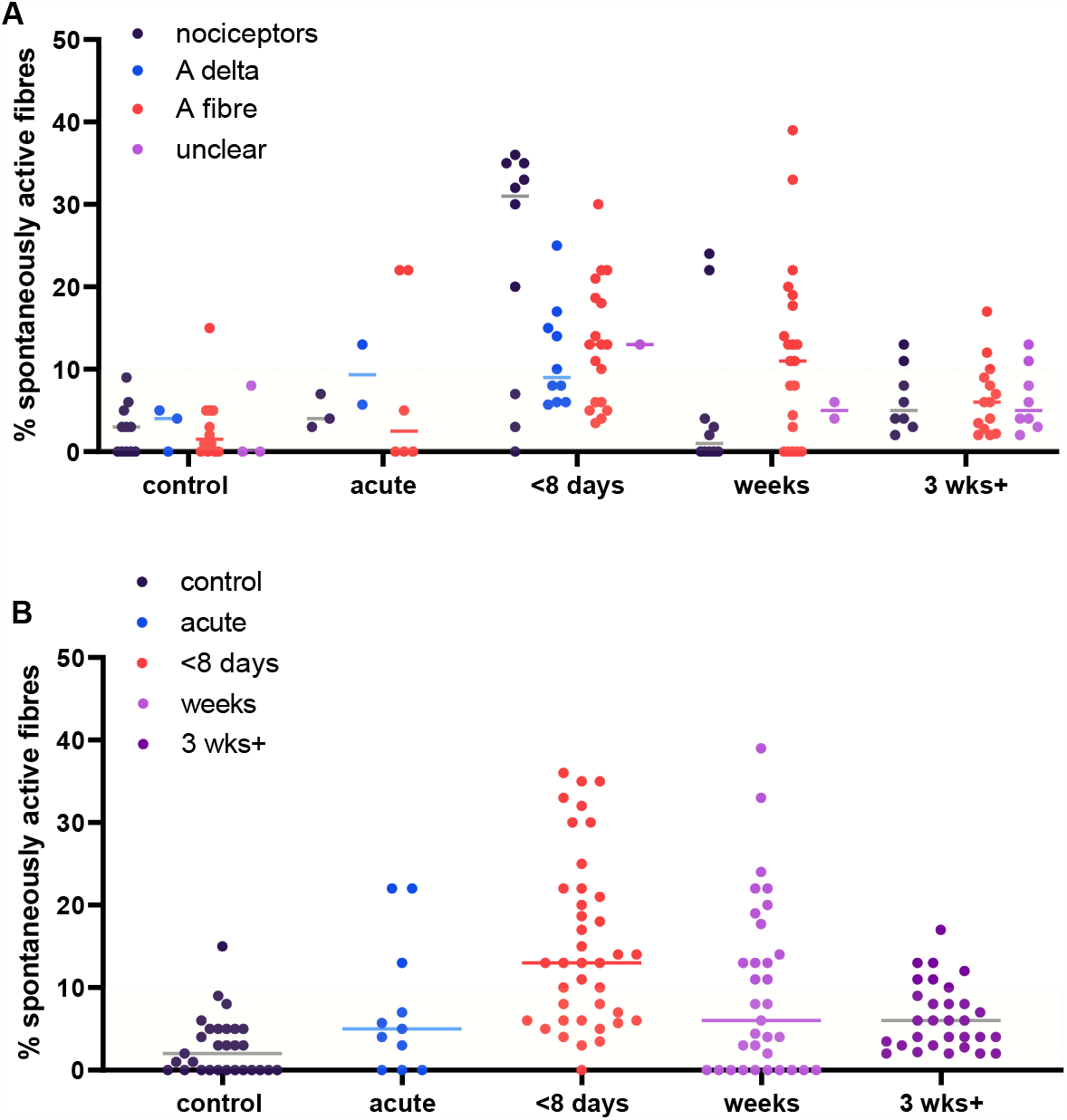
Post-injury, some spontaneous activity could be observed across all time points and fibre types, with a particularly pronounced cluster in C & Aδ nociceptors less than 8 days after injury. Each dot represents data from an experiment. Only low bias experiments were included. Injury data on mixed or cutaneous nerves, as well as from either regenerative or non-regenerative models were included. The dotted box includes experiments with less than 10% spontaneous activity, which encompassed most of the control data, derived from naïve, vehicle-treated or sham-injured animals. **A:** data split according to fibre types; **B:** fibre types pooled across time points. “control” n = 29, “acute” n = 11, “<8 days” n = 39, “weeks” n = 33, “3 weeks+” n = 30. Lines represent the median.

Examining the same time course but including only articles with their own internal controls revealed a similar picture, though it becomes clear that most data were generated less than 8 days post nerve injury (**Figure 12 & Table 4)**.

**Figure 12.**
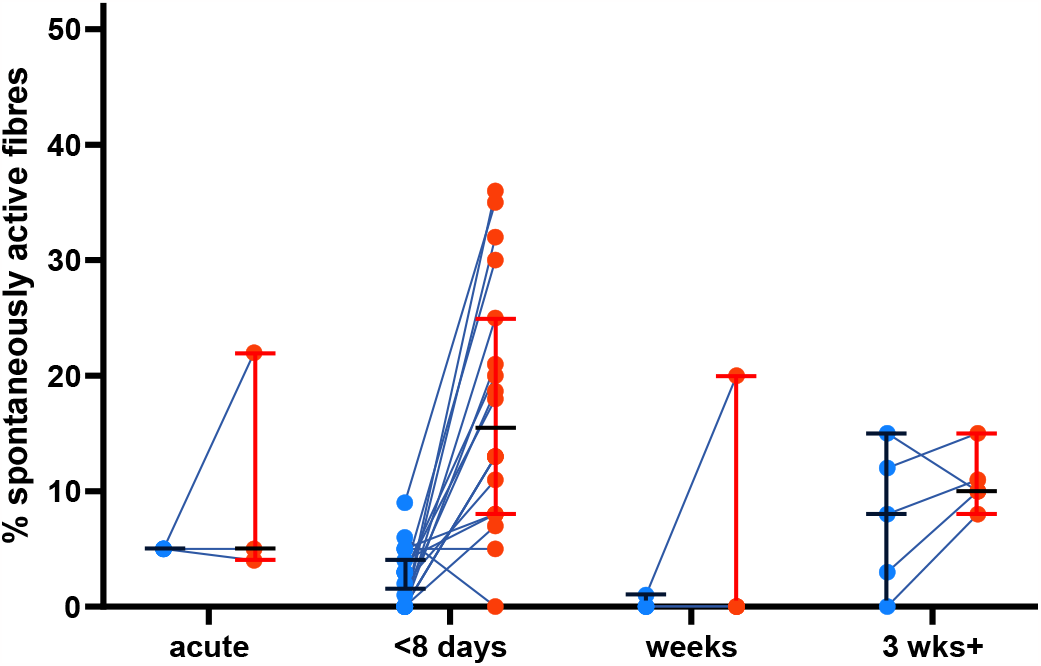
Most of the spontaneous activity data with direct control group comparisons are recorded within 8 days of injury. Each dot represents data from an experiment, lines represent medians and upper and lower quartiles. Only low bias experiments were included. Injury data on mixed, cutaneous or muscle nerves, as well as from either regenerative or non-regenerative models were included. “acute” n=3 experiments, “<8 days” n=18 experiments, “weeks” n = 4 experiments, “3 wks+” n = 5 experiments; from n = 15 studies).

## Discussion

Using our systematic search strategy and screening procedure with predefined inclusion and exclusion criteria, we present quantitative data from 147 articles performing *in vivo* electrophysiological recordings of sensory neuron spontaneous activity in animals after peripheral nerve trauma. We also collected data from 40 human microneurography experiments. Our dataset is provided in full, in an easily accessible format, in a **Supplementary Table** available [here](https://osf.io/6wpnq/?view_only=12dad701949b4ea88c2a8c3b0b50fa7c).

We found that all studies are very heterogeneous, in terms of injury type, the location of the injury, recording technique and reporting standards. Moreover, the majority of articles did not include adequate control data, with only 39% of non-human *in vivo* electrophysiology and 21% of human microneurography studies recording spontaneous activity in both non-painful and painful conditions. Finally, many datasets are small-scale in terms of sample size, and few report spontaneous activity data on a “by animal” rather than a “by fibre” basis. This obscures the variability inherent in *in vivo* electrophysiological recordings, that can be illustrated by considering some articles which do report spontaneous activity data “by animal” (**Figure 13**).

**Figure 13.**
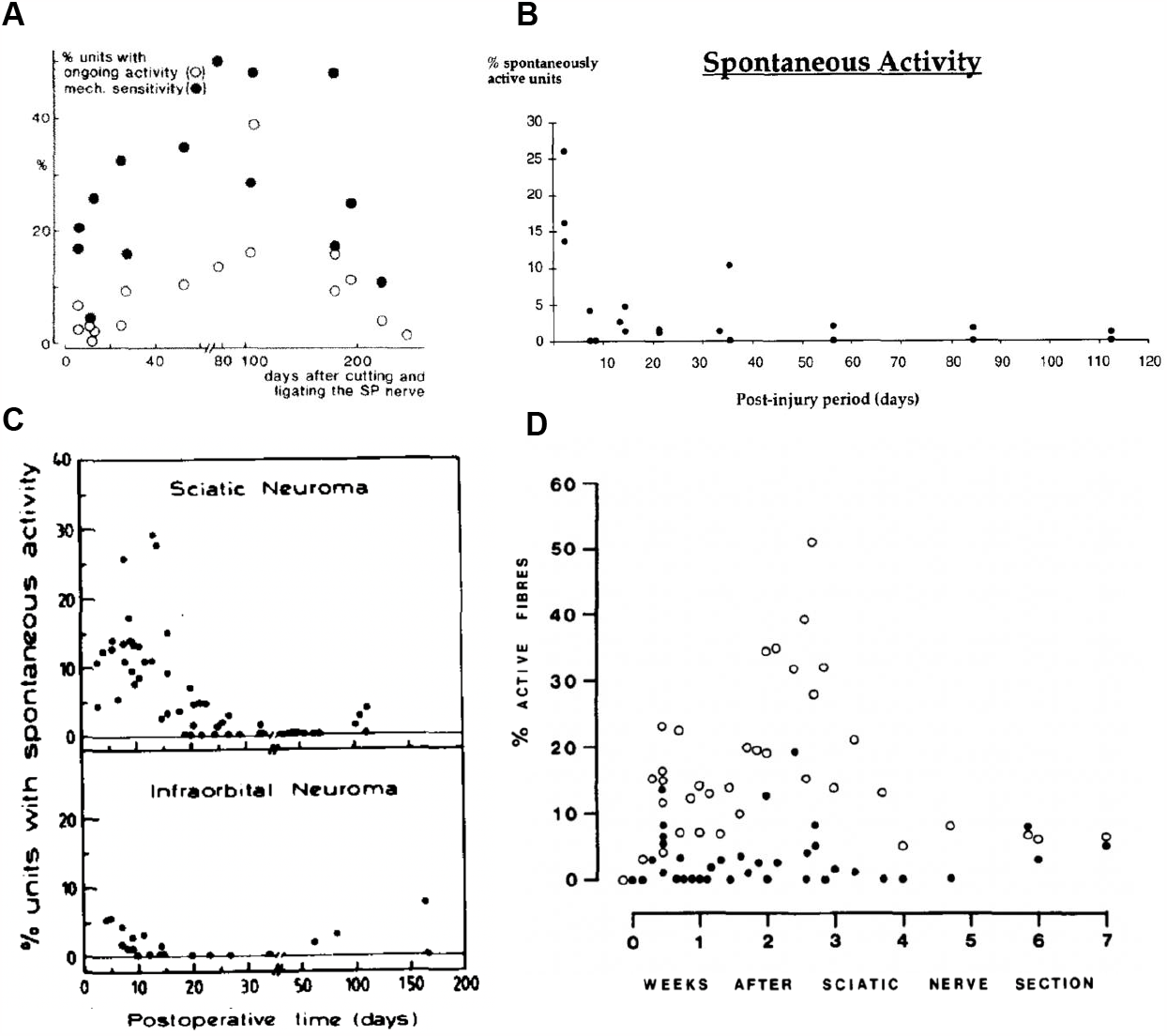
Figures derived from articles reporting animal-level data on spontaneous activity in sensory neurons at chronic time points after nerve injury. In all graphs, each dot is an animal. **A)** Transection & ligation of superficial peroneal nerves in cats to generate neuromas [3] **B)** Transection and ligation of the inferior alveolar nerve in ferrets to generate neuromas [4] **C)** Transection and ligation to generate sciatic vs. infraorbital neuromas in rats [28] **D)** Sciatic nerve transection to cause neuroma formation in mice. Closed circles denote the total percentage of spontaneous afferent activity in individual mice [23].

These studies indicate that there is very large variation between animals, even within a particular laboratory, e.g. with the percentage of spontaneously active fibres in sciatic nerve ranging from 3-30% at around two weeks after neuroma induction [28]. This highlights the serious limitation of so many articles lacking control data: without ‘internal comparators’ for individual experimental runs, it remains very difficult to estimate the true effect size of neuropathy on spontaneous peripheral neuron activity. Similarly, there is very big variation between laboratories; some of this may be due to the location of the injury, with Tal & Devor reporting that they found less spontaneous activity generated by infraorbital nerve neuromas upon direct comparison (**Figure 13C**). However, variation can still be observed when different labs report spontaneous activity after comparable injuries in comparable nerves (i.e. neuromas in the sciatic nerve or its branches, **Figure 13A, C & D**): some studies recorded spontaneous activity in about 10% of fibres within 40 days of a neuroma [3; 23], while others recorded incidence of spontaneous activity far exceeding 10% in at least half of all animals [28].

As basic scientists, we often choose to seek the explanation for such variability in biology. For example, we might argue that the differences seen between the four studies plotted in **Figure 13** are due to species differences. And yet, a more parsimonious view is that we cannot yet draw firm conclusions on biology: our review indicates that studies to date may be significantly underpowered given the large range of values that can be obtained across cohorts of animals, even within a single laboratory or an experimental paradigm. Thus, we may be able to conclude that spontaneous activity emerges in painful conditions, but a large degree of uncertainty remains around the extent and timing of this activity. This is apparent when we try and consider the questions we posed in our introduction:

### Do both A and C fibres show spontaneous activity in painful neuropathic conditions and how long does their spontaneous activity last?

Our analysis indicates that in both non-human models and in patients, painful neuropathy generally tended to induce an increase in the percentage of neurons that were being reported as spontaneously active, over and above a basal level of activity that was reported to range from 0-10% in animals and 0-4% in people. This was true across pain conditions and a range of neuropathy models, as well as A and C fibres (though in humans, only three studies examined A fibre spontaneous activity in a pain state, article numbers 95, 150 and 240 in **Supplementary Table**). It also goes some way to support the findings reported by Kleggetveit et al. [13] that people with painful neuropathy show increased spontaneous firing compared to those with non-painful neuropathy **(Figure 6**).

Animal data to date further indicate that there may be a peak time point, with spontaneous activity appearing at its maximum, particularly in nociceptive fibres, around 8 days after injury. Certainly, more than three weeks after injury, most studies reported comparatively less spontaneous activity in all types of sensory neurons. Superficially, these pre-clinical data appear in line with what has been the prevailing narrative in the pre-clinical field: A fibre spontaneous activity is more prominent and outlasts C fibre spontaneous activity [8]. Variable explanations are put forward for this phenomenon, ranging from: “a few damaged hyperexcitable nociceptors are enough to drive central sensitisation” to “a few un-damaged, hyperexcitable nociceptors are enough to drive central sensitisation” and finally “Aβ fibre spontaneous activity is what is actually driving central sensitisation”. The hypothesis implicating undamaged fibres has remained controversial, due to the difficulty of conclusively demonstrating that no fibres in the supposedly intact neighbouring spinal nerves have been accidentally damaged during complex spinal nerve transection and ligation surgeries [9].

However, it is important to note that our time series analyses are hampered by the lack of control data. Indeed, if we restrict our selection to those articles which studied specifically nociceptors more than three weeks post injury, there is only a single low-bias study (in monkey) that contains data of a non-injured control nerve [15]. Three other studies which included controls and recorded from injured fibres at such late time points either recorded from non-nociceptive A fibres [30] or did not clearly define the fibres they recorded from [7; 10]. Hence, as it stands, we feel that our systematic review data throw up a fourth possibility regarding the timing of spontaneous activity: we simply may not yet have collected sufficient, well-controlled experimental data to determine whether chronic spontaneous firing in C fibres is present in models of neuropathy. This might also be the reason for the apparent disconnect between animal and human literature: spontaneous activity in C fibres is routinely observed in patients with painful peripheral neuropathy.

### Are muscle afferents especially prone to sensitisation?

This was a view put forward as a result of direct comparison between spontaneous activity recorded from muscle versus cutaneous afferents [17]. And indeed, it is one that our systematic review data further support in an unbiased fashion. We found that the median percentage of spontaneous activity recorded in muscle nerves was significantly larger than that in mixed nerves. However, it is important to consider that studies injuring pure muscle or pure cutaneous nerves were much rarer (n = 12 each vs. n = 101 for mixed nerves). As a result, there still is a certain degree of uncertainty attached to this finding. Nevertheless, conceptually, this is potentially very important. If deep tissue afferents are indeed more prone to spontaneous activity or show a higher incidence of spontaneous activity after damage or in disease, it could explain why certain individuals with chronic pain report pain that is deep in origin [6; 18].

### Limitations

Our study had several limitations, which broadly fall into two categories: those related to systematic review methods and those related to the kind of literature we aimed to summarise.

In terms of systematic review methods, due to time constraints, we had to make some choices that deviate from best practice, even within the pre-clinical systematic review field. Specifically, we only used a single database (PubMed) for our searches, which means that we inevitably would have missed some articles; we also excluded non-English articles or those for which the full text could not be obtained easily through current subscriptions at our university; and finally, our review is not a ‘living review’, i.e. any new articles that would have appeared since May of 2022 will not be included.

Some other methodological limitations arose from the difficulties inherently linked to trying to conduct a systematic review of a very heterogeneous set of pre-clinical literature. The considerable challenge associated with designing search strings and data extraction strategies for such a diverse set of articles meant that we had to deviate quite significantly from the protocol we originally registered. Specifically, we had to design a whole new search string when we realised that our first one was too limited, with certain MeSH terms filtering out key articles in the area we aimed to cover. Moreover, we initially were much too ambitious in our data extraction strategy, and ultimately had to limit the data we derived to make the workload manageable for two people within the time frame of this research project. For this last reason, we focused our analysis on *in vivo* electrophysiological studies and excluded studies on spontaneous activity in inflammatory states. The latter is arguably not a bad choice from a conceptual point of view, since neuronal firing in these models is less likely to be ‘truly spontaneous’ [2].

Beyond methodological constraints, our dataset is limited by the nature of the articles we reviewed. For example, we were unable to summarise our data in more conventional review formats, e.g. forest plots of effect sizes. This was mainly due to the majority of studies not containing control groups, making it impossible to calculate the relative increase in spontaneous activity between case & control for many experiments. We also had to contend with the fact that most studies record their results on a ‘by-fibre’ rather than a ‘by-animal’ basis. As a result, animal sample sizes in this area are often not very informative. Finally, given the historical nature of much of the literature we summarised, there was almost no standardisation of recording setups, recording protocols, or reporting standards.

## Conclusions

We aimed to use systematic search and data extraction methods to shed light on what is known about spontaneous activity in peripheral sensory nerves from *in vivo* electrophysiology and microneurography studies. We conclude that the literature suggests that spontaneous activity is regularly detected in both non-nociceptive A fibres and nociceptors after injury in animals and in human patients with ongoing neuropathic pain. There also appeared to be indications that spontaneous activity was more prominent in muscle afferents and in painful rather than non-painful neuropathy.

Our review also highlights surprising uncertainty in our knowledge beyond these very basic statements. Due to highly heterogeneous recording protocols, lack of controls or reports on inter-animal variation, we were unable to estimate effect sizes or make strong statements regarding the emergence and duration of spontaneous activity in different fibre types.

Moreover, our data suggest that many apparent disconnects between human and rodent recordings may simply be due to comparing models and timelines that are not equivalent. For example, 76% of all rodent work is conducted on non-regenerating traumatic nerve injury, while there is only a single microneurography study that might arguably be equivalent (on amputees who might or might not suffer from neuromas). Similarly, only a single well-controlled animal study exists that examines specifically C-fibre spontaneous activity more than three weeks after nerve injury with *in vivo* electrophysiology. Indeed, when humans and rodents were directly compared using microneurography, species differences appear to diminish [25; 26]. The incidence of spontaneous activity in all 1134 nociceptors recorded in various rat models of neuropathy was found to be at 19.8%, while in humans with small fibre neuropathy, out of 102 fibres, 20.6% were spontaneously active [26]. Moreover, just like in human, microneurography could identify spontaneously active C fibres in rats many months after nerve injury, with the longest time point assessed being 287 days post-surgery [25].

We therefore must end with a very clear plea for the generation of more, and more standardized, electrophysiological data in this area. Novel, higher-throughput proxy-measures of sensory neuron activity, such as *in vivo calcium* imaging, could also be employed [12]. While we recognise that these types of experiments are extremely technically demanding and time-consuming, they are also vital for us to understand perhaps one of the most immediate causes of chronic pain – that of ongoing and/or spontaneous activity in peripheral sensory neurons.

## Acknowledgements

DC was funded by a PhD studentship from Mundipharma. FD is the recipient of a Medical Research Foundation Prize (MRF-160-0015-ELP-DENK-C0844). GG is funded by an *Advanced* Pain Discovery Platform UKRI MRC grant *(MR/W027518*/1). BN is supported by the German research council (DFG NA 970 3-1, 6-2 and 7-2 and supported by a grant from the Interdisciplinary Center for Clinical Research within the faculty of Medicine at the RWTH Aachen University. NS is funded by a Jennie Gwynn post-doctoral career development fellowship. None of the authors have any conflicts of interest to declare.

We would like to thank and commemorate Stephen McMahon, who was involved in the early stages of this project, but sadly passed away in late 2021.

Figures were partly generated using Servier Medical Art, provided by Servier, licensed under a Creative Commons Attribution 3.0 unported license.

This research was funded in whole or in part by the UK Research & Innovation. For the purpose of Open Access, the author has applied a CC BY public copyright licence to any Author Accepted Manuscript (AAM) version arising from this submission.

